# ToxiVerse: A Public Platform for Chemical Toxicity Data Sharing and Customizable Predictive Modeling

**DOI:** 10.64898/2026.02.26.708255

**Authors:** Prasannavenkatesh Durai, Daniel P. Russo, Yitao Shen, Tong Wang, Elena Chung, Lang Li, Hao Zhu

## Abstract

Chemical toxicity assessment is critical for drug development and environmental safety. Computational models have emerged as a promising alternative to animal testing and now play a significant role in efficiently evaluating new chemicals. To address the urgent need for providing user-friendly machine learning tools in computational toxicology, we developed ToxiVerse, a public web-based platform. It provides curated toxicity datasets, automatic chemical bioprofiling, and a predictive modeling interface designed for researchers who lack programming expertise. The platform comprises three integrated modules: (i) the Bioprofiler module, which provides chemical descriptors by combining chemical-bioactivity data from PubChem assay with a machine learning-based data gap-filling procedure; (ii) the Database module, which hosts around 50,000 curated unique chemicals covering diverse toxicity endpoints; and (iii) the Cheminformatics module, which allows users to upload their own datasets, use datasets from ToxiVerse, or retrieve existing data from PubChem; perform chemical curation; and automatically generate Quantitative Structure-Activity Relationship (QSAR) models to predict chemicals of interest. ToxiVerse enables researchers to carry out bioprofiling, access curated toxicity datasets, and evaluate chemical toxicity through machine learning-based modeling and prediction. The platform is supported by sample files and a detailed tutorial, and it is freely accessible at www.toxiverse.com.

**GRAPHICAL ABSTRACT:** 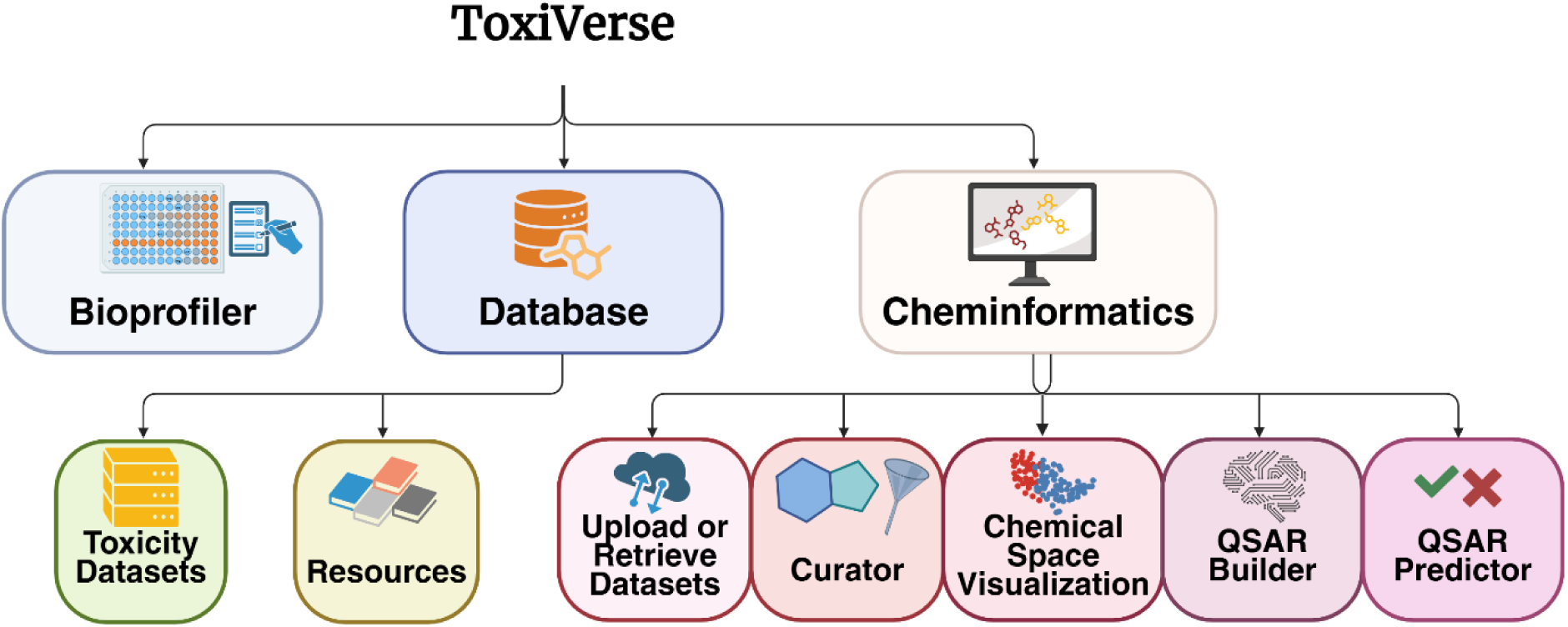

## INTRODUCTION

Chemical safety evaluation is fundamental to safeguarding both human and environmental health and underpins decisions made by regulatory agencies (1). In drug development, drug-induced side effects and toxicities are major contributors to both early- and late-stage failures, responsible for up to 40% of preclinical attritions reported in 2015 and about 20% of Phase II and Phase III attritions reported in 2008 (2,3). Chemical toxicity also imposes a major public health burden, with more than two million adverse drug events and approximately 100,000 deaths reported in the United States in 1994 alone, linked to drug-induced toxicity (2). Current regulatory frameworks still rely heavily on animal testing, which is time-consuming, costly, and ethically controversial (4–6). These limitations have catalyzed a shift toward alternative approaches, including *in vitro* testing and *in silico* modeling (4). The progress of computational toxicology along with this shift has been driven by advances in computational power and new algorithm development, as well as the increasing availability of large public databases of toxicity data (7–9). For instance, Tox21 (10) has screened more than 10,000 compounds across diverse *in vitro* assays to support high-throughput, mechanism-informed toxicological profiling. The resulting toxicity data are publicly available through the Tripod database (https://tripod.nih.gov/tox/) and NIEHS Integrated Chemical Environment (https://ice.ntp.niehs.nih.gov/DATASETDESCRIPTION?section=cHTS) (10).

Building on the increasing availability of data and new algorithms, various computational tools have been developed to meet the growing demand for new chemical toxicity evaluations (11). These include online platforms such as ADMETlab 3.0 (12), admetSAR 3.0 (13), ProTox 3.0 (14), EMolTox (15), and VenomPred 2.0 (16), as well as tools for specific toxicities like Pred-hERG 5.0 (17), PredSkin 3.0 (18), and CardPred (19). For example, ADMETlab 3.0 is a web-based platform that can predict over 80 endpoints related to Absorption, Distribution, Metabolism, Excretion, and Toxicity (ADMET) using Directed Message Passing Neural Network (DMPNN) models (12). The admetSAR 3.0 tool supports predictions for over 100 ADMET endpoints and combines classification and regression models built with a contrastive learning-based multi-task graph neural network framework (CLMGraph) (13). ProTox 3.0 integrates chemical structures, protein targets, and pathway information as outputs and predicts 28 toxicity endpoints using models developed with deep neural network (DNN) or Random Forest (RF) methods (14). Other toolboxes like QsarDB (20) offer a digital repository of validated Quantitative Structure-Activity Relationship (QSAR) models, while VEGA-QSAR (21) and the OECD QSAR Toolbox provide built-in models and read-across tools for predicting chemical toxicities (22).

Recent improvements in these chemical toxicity evaluation tools include integrating deep learning methods (e.g. DNN, DMPNN, and CLMGraph) into model developments, which enhance model performance and broaden endpoint coverage (7). However, model interpretability continues to be a challenge, particularly in deep learning-based models due to their black-box nature (9,11). Training data size and quality also impacts model predictivity. Many publicly available toxicity databases suffer from issues such as inconsistent annotations, missing metadata, lack of rigorous curation, non-standardized data formats, and species discrepancies (7,23). These problems can introduce significant bias into toxicity modeling results. Few existing chemical toxicity evaluation tools offer essential features such as batch processing, support for custom dataset modeling, and standardized modeling reports (11). Moreover, most existing computational toxicity tools require programming expertise, making them less accessible for researchers without such backgrounds (7,11). Although web-based platforms are popular, many rely only on pre-trained models and offer limited flexibility for user data input and customized predictions (7,11,24).

In this study, we present ToxiVerse, an online chemical toxicity modeling platform designed to address these challenges by delivering a user-friendly web-based toolbox that integrates automated chemical bioprofiling based on PubChem assay data, curated toxicity datasets across diverse endpoints, and customizable machine learning (ML) modeling. ToxiVerse has three application modules. First, the Bioprofiler module that profiles chemicals against their bioassay outcomes from PubChem (25–28) and predicts outcomes for target chemicals with inconclusive or missing data, thereby generating comprehensive chemical descriptors. Second, the Database module provides access to rigorously curated chemical toxicity datasets covering around 50,000 chemicals, mostly with data on critical endpoints such as hepatotoxicity, carcinogenicity, and developmental toxicity. Third, the Cheminformatics module offers utilities for dataset upload or retrieval from the PubChem database, dataset curation, and automated QSAR model generation using widely adopted ML algorithms. Users can build models using any provided ToxiVerse datasets or their own data and select the most suitable models to predict their chemicals of interest. By providing a user-friendly ML platform, ToxiVerse empowers researchers worldwide to perform interpretable, scalable, and computational toxicity modeling for chemical risk assessment purposes.

## MATERIALS AND METHODS

### Platform implementation

ToxiVerse platform was developed as a modular web application using the Flask framework (Flask 2.2, Python 3.11) and deployed within a Docker environment. The front-end interface was implemented with HTML and CSS, JavaScript and jQuery 3.4.1, styled using Bootstrap 4.1.0. Interactive visualizations are implemented with Plotly 5.9.0, while server-side plots are rendered using Matplotlib 3.7.0 and Seaborn 0.13.0. Server-side execution is handled by Gunicorn, while Redis 4.4.0 is employed for asynchronous task queuing via Redis Queue. Curated datasets and PubChem chemical-bioactivity data are stored in SQLite database, accessed through SQLAlchemy 3.0.2. A RESTful API supports core functionalities, including Principal Component Analysis (PCA) visualization, chemical distribution plotting, access to relevant bioassay data of endpoints, and dataset downloads. Data curation process and descriptor calculation were performed using RDKit 2022.09.2 (www.rdkit.org), while predictive modeling was carried out with scikit-learn (1.2.0). The web server is publicly accessible without requiring user login.

### Bioprofiler module

#### Database Construction with PubChem Data

A SQLite database leveraging PubChem data was built using two primary datasets downloaded from the PubChem FTP (29) (accessed on April 10, 2025): a bioactivities set and a compound-to-InChIKey mapping set. The bioactivities set contains experimental assay results, including PubChem compound identifiers (CIDs), PubChem assay identifiers (AIDs), and reported chemical activity outcomes (*active*, *probe*, *inactive*, *inconclusive, or unspecified*). The mapping set provides standardized chemical structures in the form of InChI strings and InChI Keys for each CID. These two datasets were merged based on CID, and records with missing or incomplete InChI strings were removed to ensure consistent chemical representation. The final database used for profiling in this study contains five key fields: CID, AID, Activity Outcome, InChI string, and InChI Key.

#### Assay Selection and Machine Learning Model Development

Mutual information (MI) scores were calculated using scikit-learn’s classification module to guide assay selection. MI measures how informative assay outcomes are about the overall activity of chemicals. First, the bioactivity data collected from PubChem assays for target chemicals were transformed into a chemical-bioactivity matrix (i.e. initial bioprofile), where active values were encoded as 1 and inactive or inconclusive values were encoded as 0. Next, each chemical was assigned a binary overall activity label, defined as active if it was active in at least one assay. Finally, the MI between the activity outcome of each assay and the overall activity label was calculated. An assay achieves a high MI score when both its actives (1) and inactives (0) closely match the overall activity labels of as many chemicals as possible. The more consistently an assay distinguishes overall active from inactive chemicals, the higher its MI is.

For chemicals from the selected assays, ML models were developed to fill data gaps for compounds lacking experimental *active*/*inactive* confirmation. To this end, RF classifiers were trained using Extended-connectivity fingerprints (ECFP) 6 with a radius of 3 and a bit length of 2048, and their performance was evaluated using commonly adopted binary classification metrics. RF is an ensemble learning algorithm that constructs many individual decision trees during training and aggregates their outputs to generate predictions (30). ECFP are circular topological fingerprints that encode local atomic neighborhoods into fixed-length binary vectors (31). In this function, ECFP6 fingerprints are generated using a bond radius of three, producing 2048-bit vectors that capture substructural patterns up to six bonds in diameter.

### Database module

Curated chemical datasets were collected from multiple resources, most of which have been described in our previous studies (32–45). These datasets were used to construct an SQLite database, where each chemical was assigned a unique ToxiVerse-ID, canonical SMILES were generated and stored, and PubChem CID was included when available. Each dataset can be downloaded and explored through a histogram showing the distribution of chemical counts across activity value ranges.

In addition, relevant bioassays for each endpoint across all datasets were extracted using bioactivity data collected from PubChem with the Bioprofiler module. Based on the bioprofile data of chemicals in each dataset, bioassays relevant to the corresponding endpoint were identified. First, the counts of *active*, *inactive*, and *inconclusive* compounds were computed for each bioassay in the bioprofile data. The active rate for bioassay *i* was calculated as:

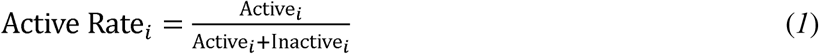

To harmonize active rates while accounting for the active rates of other assays in the bioprofile, Bayesian smoothing was applied to adjust the active rates calculated above. The average active rate μ across all relevant bioassays was computed as:

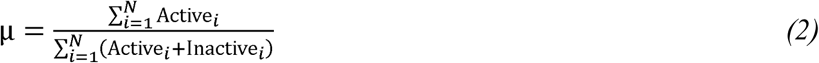

where *N* denotes the total number of bioassays in the bioprofile. For each bioassay *i*, the Bayesian Adjusted Score was then calculated as:

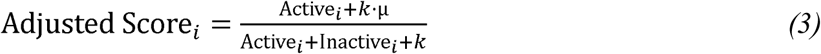

where *k* = 100 is the Bayesian smoothing parameter that balances the influence of the average activity rate with assay-specific observations. The calculated Adjusted Score was then used to rank the bioassays for their relationships to the target endpoint.

### Cheminformatics module

#### Dataset and Chemical Curator

Input chemical files, in either CSV or SDF format, can be uploaded by users through the Cheminformatics module. Users can also import datasets directly from PubChem by providing the relevant AID as input. The curator function leverages scripts from the ChEMBL Structure Pipeline (46), an open-source cheminformatics framework designed to systematically clean, standardize, and validate the input chemical structures. Built on top of RDKit, the pipeline performs rigorous chemical structure curation to ensure chemical integrity and consistency. This procedure detects invalid or ambiguous stereochemistry, problematic valence states, illegal bond types, and undesirable substructures (e.g., salts, solvents, metals). Then the standardization step includes kekulization, hydrogen removal, SMIRKS-based normalization, uncharging, and conformer cleanup.

#### Chemical space visualization

The PCA dimensionality reduction method was employed to visualize the chemical space of the datasets. Descriptors are calculated using RDKit and then standardized. Dimensionality reduction is performed with scikit-learn to extract the top three principal components. The resulting PCA coordinates, together with CIDs and activity labels, are used to generate a 3D scatter plot. Compounds are color-coded by activity classification, enabling users to explore chemical clusters within each dataset.

#### QSAR Modeling

Based on user selection, molecular descriptors or fingerprints (RDKit molecular descriptors, ECFP6 fingerprints, and Functional-Class Fingerprint (FCFP) 6) are calculated for each chemical in the target dataset and normalized if necessary. FCFP are feature-based circular fingerprints that encode local pharmacophoric environments into fixed-length binary vectors (31). In this function, FCFP6 fingerprints are generated using a bond radius of three, resulting in 2048-bit vectors that capture feature-level substructural patterns up to six bonds in diameter. The module supports classification or regression modeling using ML algorithms, including RF, Support Vector Machines (SVM), and k-Nearest Neighbors (k-NN). Model hyperparameters are optimized via grid search with five-fold cross-validation. For classification, a default probability threshold of 0.5 is used to classify chemicals (1 = active/toxic, 0 = inactive/non-toxic). The best-performing model by cross-validation metrics can be saved for future predictions. SVM is a supervised learning algorithm that identify the optimal separating hyperplane between classes in a high-dimensional feature space (47). The *k*-NN is a distance-based, non-parametric algorithm that classifies a query compound according to the majority class of its k most similar compounds in the training set (48).

#### QSAR Prediction

By selecting a generated model, chemicals to be predicted can be input through two options: (i) file upload as a CSV or SDF file, or (ii) direct entry of SMILES strings. For CSV files, the system automatically detects a column named “SMILES” or allows users to manually specify the column containing the SMILES strings of the chemicals. The predicted results are provided as CSV or SDF files.

## RESULTS AND DISCUSSION

### Overview of ToxiVerse

ToxiVerse is a web-based informatics platform designed to provide researchers with a bioprofiling tool for target chemicals, curated chemical toxicity datasets, and computational toxicity modeling protocols within a user-friendly working environment. The platform includes a Tutorial page with detailed instructions for efficient use. The three modules of ToxiVerse and their functions are shown in Figure 1.

**Figure 1.**
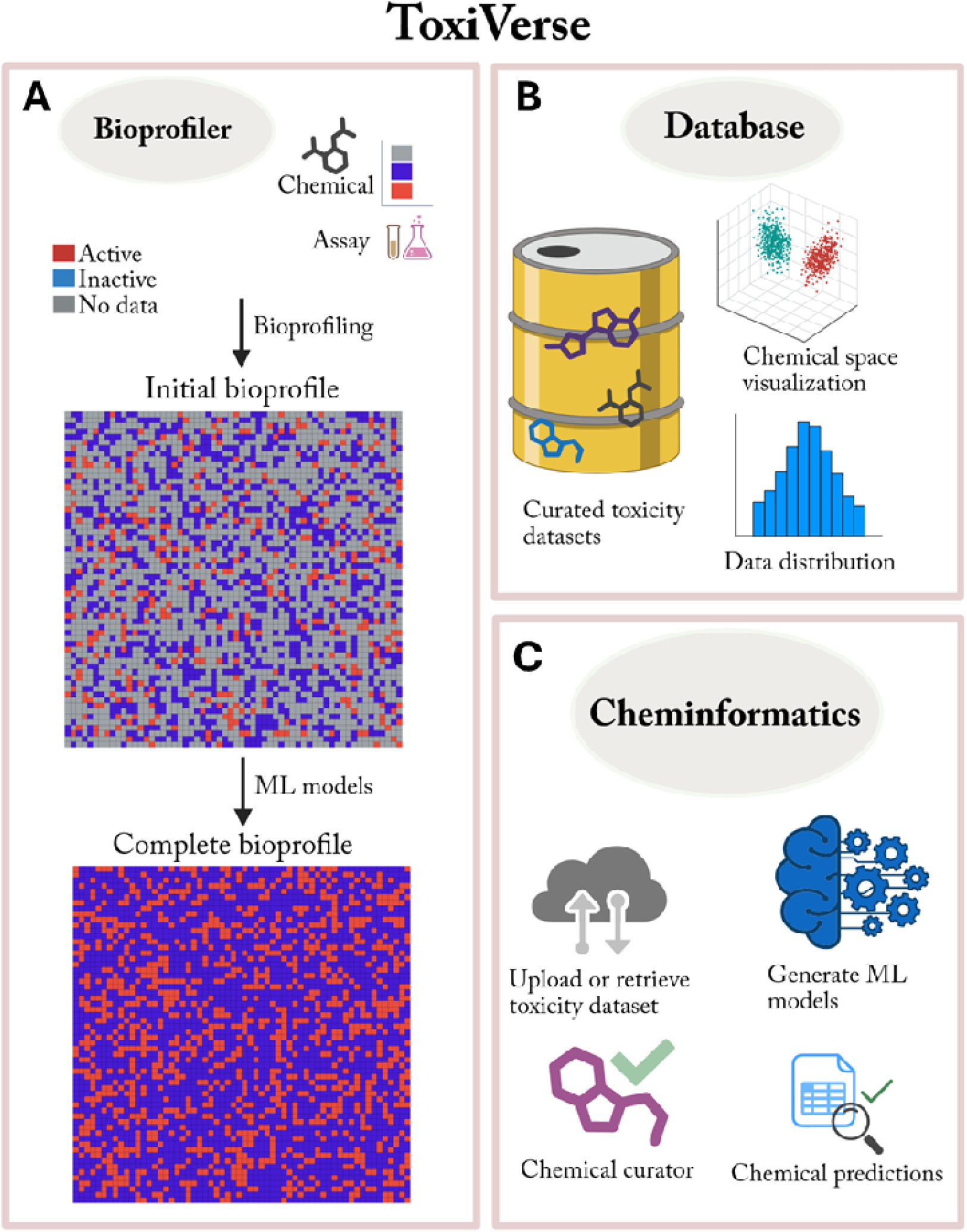
Overview of the three main modules in ToxiVerse. (**A**) Bioprofiler (**B**) Database, and (**C**) Cheminformatics. The functions available within each module are shown.

The Bioprofiler module constructs initial bioprofiles for chemicals by leveraging bioassay data from PubChem and provides complete bioprofiles after data gaps are filled with ML models. It allows users to input chemicals and generates a chemical-bioactivity matrix (i.e., an initial bioprofile), which can be used to identify key assays relevant to the chemicals of interest (i.e., assays that distinguish between Actives/Toxics and Inactives/Non-toxics). ML models built from chemicals in selected assays are then used to impute prediction values for input chemicals lacking experimental data in these assays. The Database module hosts a curated collection of approximately 50,000 chemicals, mainly covering toxicological datasets along with detailed data information. The module includes functions to download selected datasets and access PubChem bioassays relevant to specific toxicological endpoints. The Cheminformatics module provides options for dataset upload or retrieval from PubChem, dataset curation, visualization, and QSAR modeling and prediction. Users can upload their own datasets, use the ToxiVerse datasets provided, or import data directly from PubChem for modeling and prediction purposes.

### Bioprofiler

The Bioprofiler module enriches the bioactivity of target chemicals by integrating PubChem assay outcomes with model-predictions for untested assays. This approach provides bioactivity-based chemical descriptors that embed compounds in a biologically informed feature space that extends beyond chemical structure alone. High-throughput screening (HTS) assays in PubChem form the basis of this approach by revealing how chemicals interact with molecular targets and indicating potential toxicity pathways (29). To create biologically enriched data for target chemicals, the Bioprofiler first constructs a chemical-bioactivity matrix (initial bioprofile). MI scoring is then applied to select the most informative assays for downstream modeling.

Assay-specific RF models are used to impute missing outcomes, producing a more complete bioprofile for each compound. This gap-filling strategy enables chemical read-across and ensures that target compounds are represented with comprehensive bioactivity profiles. A key advantage of this activity-based profiling is the ability to achieve mechanistic related biological data for target chemicals. Compounds with little structural resemblance may nevertheless display similar activity across assays, reflecting shared mechanisms of action or phenotypic effects. By combining observed and predicted assay outcomes, the resulting hybrid descriptors integrate structural and biological information. This enriched potential training data can improve comprehensive toxicity modeling compared with structure-only descriptors. Overall, the Bioprofiler module transforms sparse HTS data into comprehensive bioactivity descriptors for target compounds. This maximizes the value of public big data and supports computational toxicology, risk assessment, and read-across applications. The integration of chemical-bioactivity profiling with statistical analysis and data gap filling has been validated across multiple toxicity prediction studies, demonstrating its reliability and effectiveness. (36,49–51).

#### PubChem Bioassay Data

To facilitate efficient retrieval of relevant bioactivity data and other useful information (e.g., chemical identifiers) for user-input chemicals, an SQLite database was generated from PubChem data containing chemicals and their assay outcomes. The database is updated every six months to remain consistent with current PubChem releases. The resulting database integrates assay bioactivities with standardized unique chemical identifiers and structural representations, enabling rapid querying without repeated network inquiries to PubChem servers, which would otherwise require significant time. The current SQLite database includes CIDs, AIDs, activity outcomes, InChI strings, and InChIKeys collected from PubChem.

#### Bioprofiling

The core functions of the Bioprofiler module are shown in Figure 1A. Bioprofiler module supports automatic bioassay data retrieval from local PubChem database for input chemicals. Bioactivity data are collected from PubChem assays that have historically tested the input chemicals. The initial bioprofile contains AIDs (assay identifiers), CIDs (chemical identifiers), and activity outcomes. Activity outcomes are encoded as 1 for Actives/Toxics, -1 for Inactives/Non-toxics, and 0 for Inconclusive/Unspecified/No data. When different results exist for the same CID-AID pair, the active value is retained.

Figure 2 displays the Bioprofiler interface for file upload and results download. The initial bioprofile is a high-dimensional matrix with chemicals as rows and assays as columns. Only assays with at least one active compound are retained. Users can download this matrix via the *Bioprofile* option and a heatmap representation via the *Heatmap* option. A clustered heatmap generated from 1622 PubChem assays for an estrogen receptor toxicity dataset of 6543 compounds (45) is shown in Figure 3.

**Figure 2.**
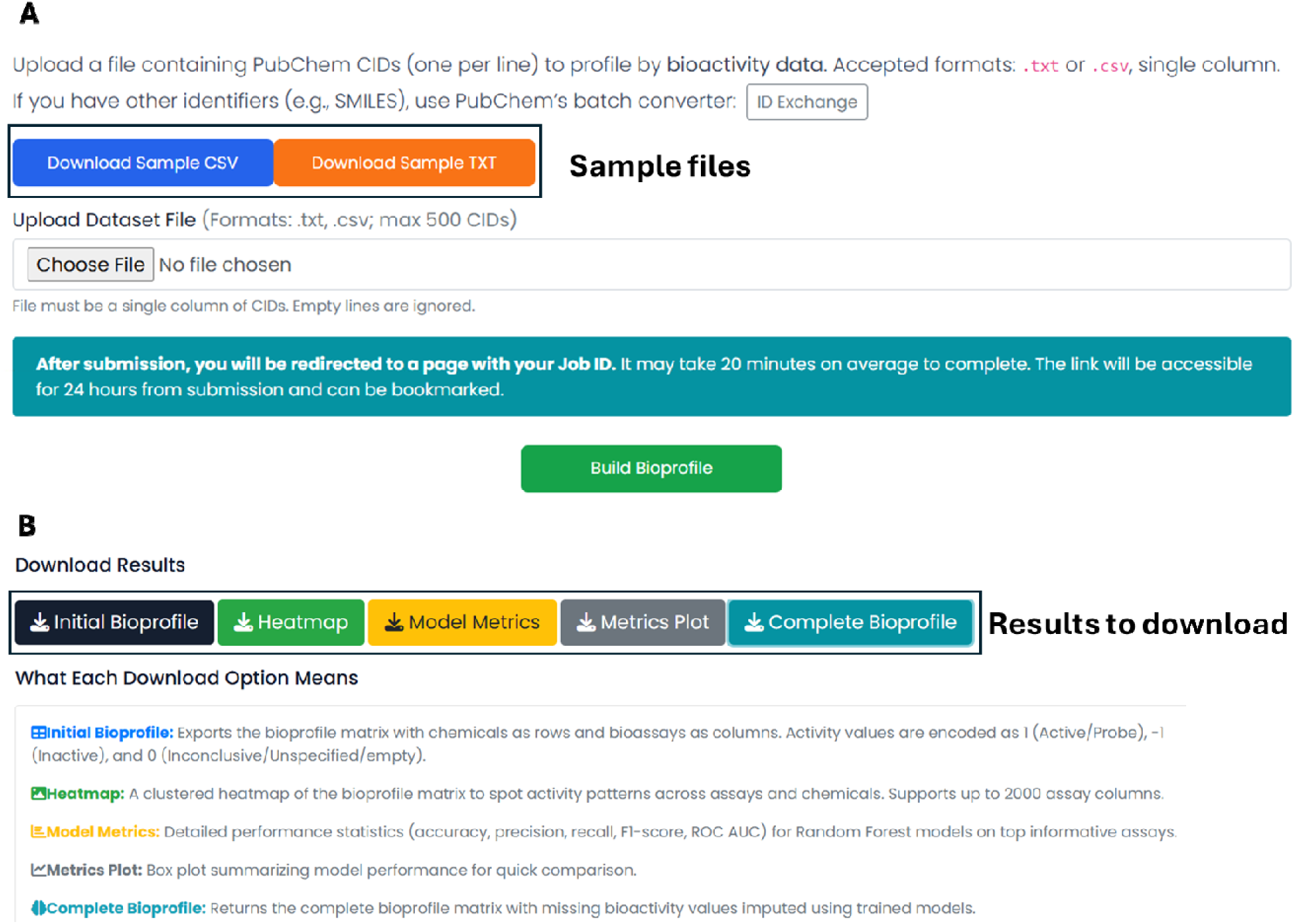
The Bioprofiler interface shows (A) input options for file upload and (B) result download options along with result interpretation.

**Figure 3.**
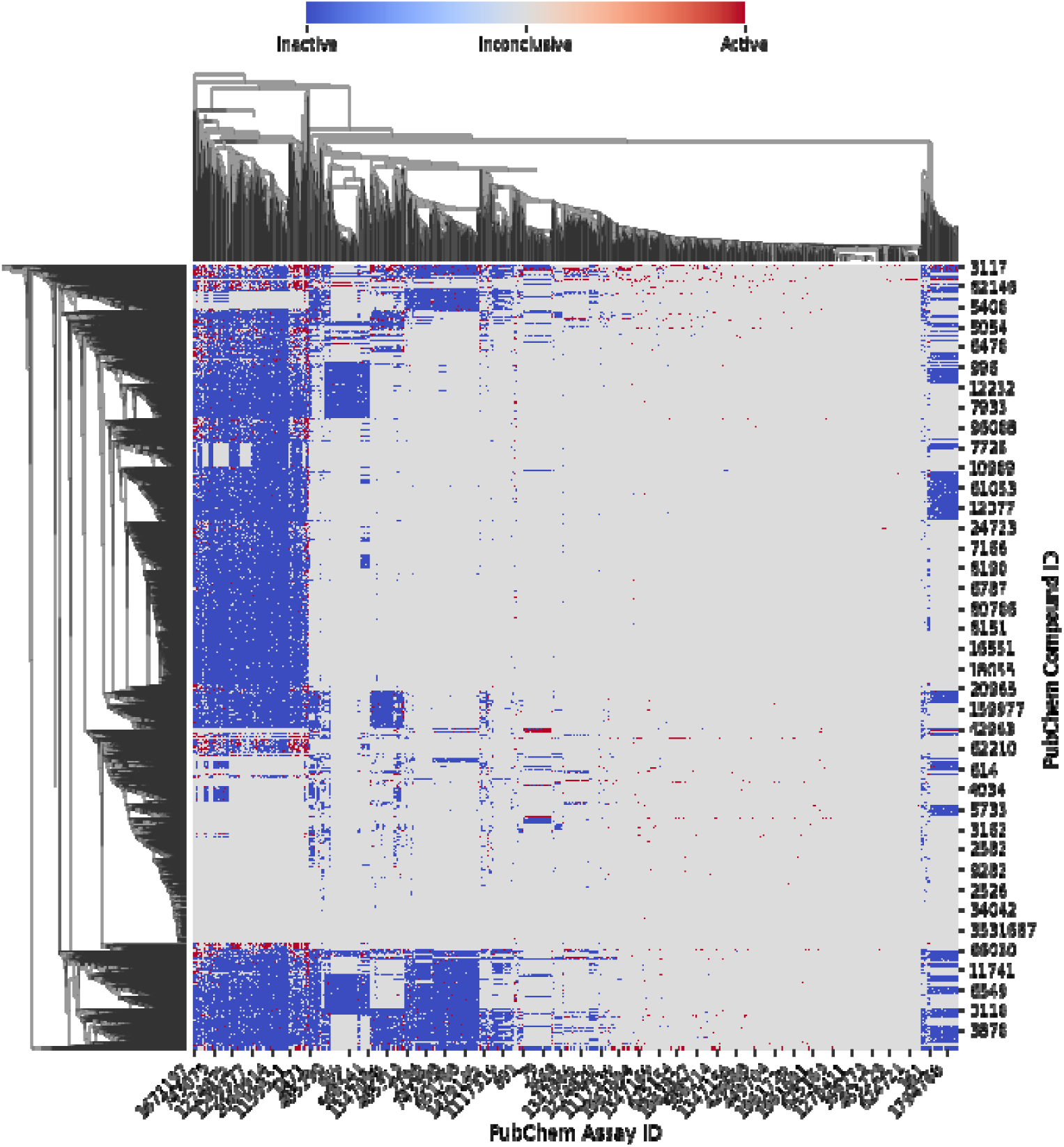
Heatmap of the bioprofile for an estrogen receptor toxicity dataset. Red = active (1), blue = inactive (-1), grey = inconclusive (0).

MI scores are computed to rank assays for their usefulness in data analysis and modeling. A high MI score indicates an assay with sufficient actives that can be used to explain potential mechanisms of target toxicity, making it highly informative for modeling. Since target chemicals have missing experimental data against many PubChem assays, data gaps in the initial bioprofile are filled using chemicals originally tested in these assays. For each selected assay, up to 500 actives and 500 inactives are randomly selected from the chemicals being tested, and only assays with at least 100 chemicals in both classes are retained to ensure resulting model predictivity. Binary RF models are trained using ECFP6 fingerprints generated in RDKit. Model evaluation files are available via the *Model Metrics* option, and plots can be obtained via the *Metrics Plot* option. The trained models are then used to predict assay outcomes for target chemicals lacking experimental data (previously encoded as 0), producing a complete bioprofile downloadable via the *Complete Bioprofile* option. The results can be accessed for up to 24 hours from job submission via unique job link. The results of entire Bioprofiler module are presented in Supplementary Figure S1. The performance metrics from the acute toxicity dataset (23) are shown as a box plot in Supplementary Figure S1B. For input chemicals with CIDs, five downloadable outputs are available (Figure 2B).

### Database

The *Database* module provides curated datasets that are standardized, user-friendly to access, and ready for modeling. High-quality curated datasets are critical for advancing computational toxicology (7,11). The database currently contains approximately 50,000 chemicals drawn primarily from our previous studies (32–45), complemented by additional toxicity data from public resources. These datasets mainly cover toxicity data spanning more than 50 endpoints, including widely studied areas such as acute toxicity, hepatotoxicity, endocrine disruption. (Figure 4). Each dataset only consists of unique chemicals linked to well-known identifiers (SMILES, and PubChem CID) along with ToxiVerse ID, making them suitable cheminformatics modeling pipelines.

**Figure 4.**
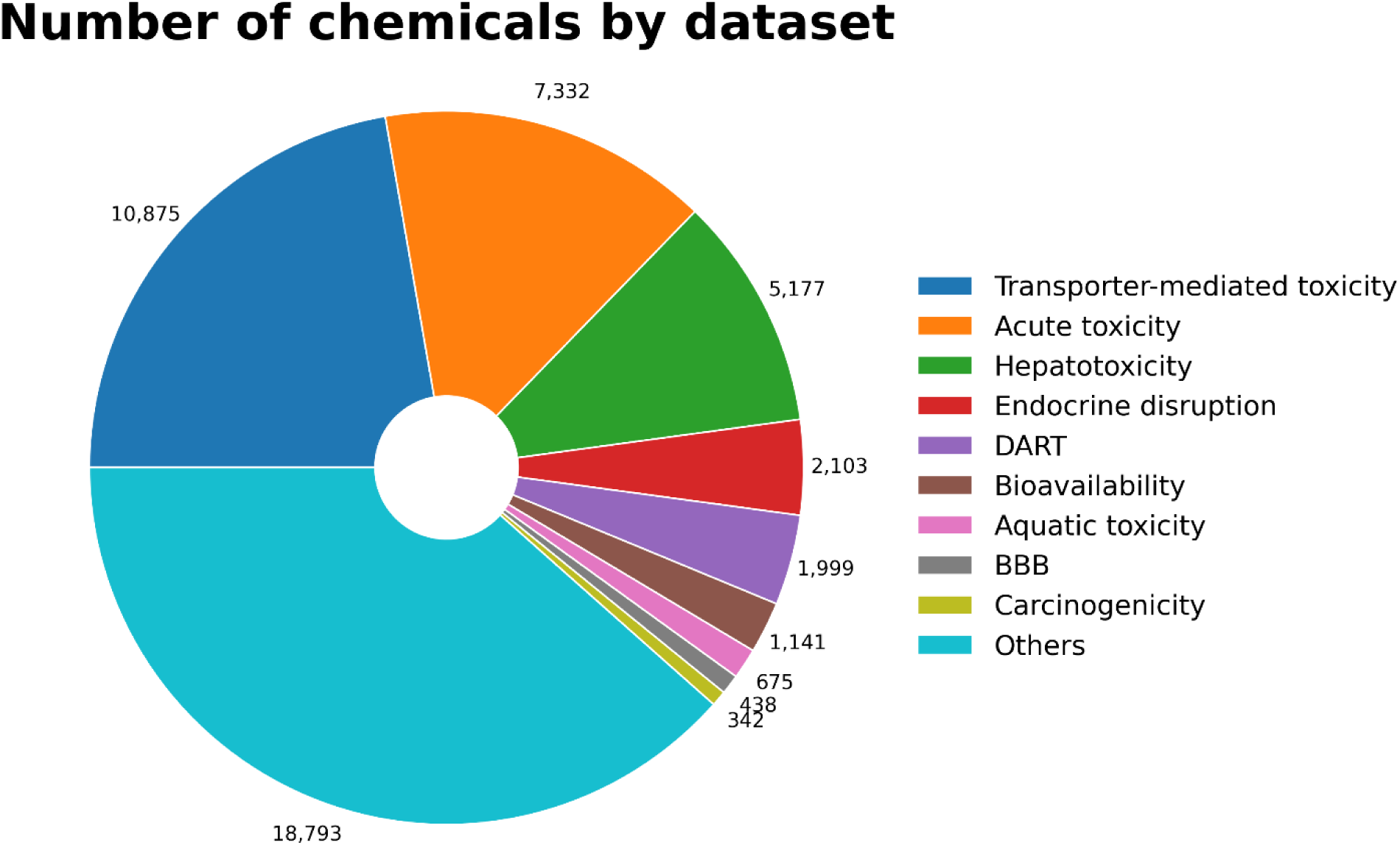
Number of chemicals in each dataset by category.

Data accessibility of this platform is easy and user friendly. Users can interactively explore chemical space, view endpoint distributions, and download curated CSV files with a single click. By consolidating multiple toxicity resources into a single harmonized framework,

ToxiVerse overcomes common challenges of data availability and usability, ultimately promoting broader adoption of computational approaches in toxicology.

When a toxicity endpoint in a dataset is selected, users have three integrated views of the relevant data (Figure 5): a PCA plot that shows the chemical space distribution of the chemicals in the dataset, a histogram illustrating the distribution of activity/toxicity values across chemicals, and a list of the 500 PubChem assays most relevant to the endpoint, identified using the Bayesian scoring approach (Equations 2&3).

**Figure 5.**
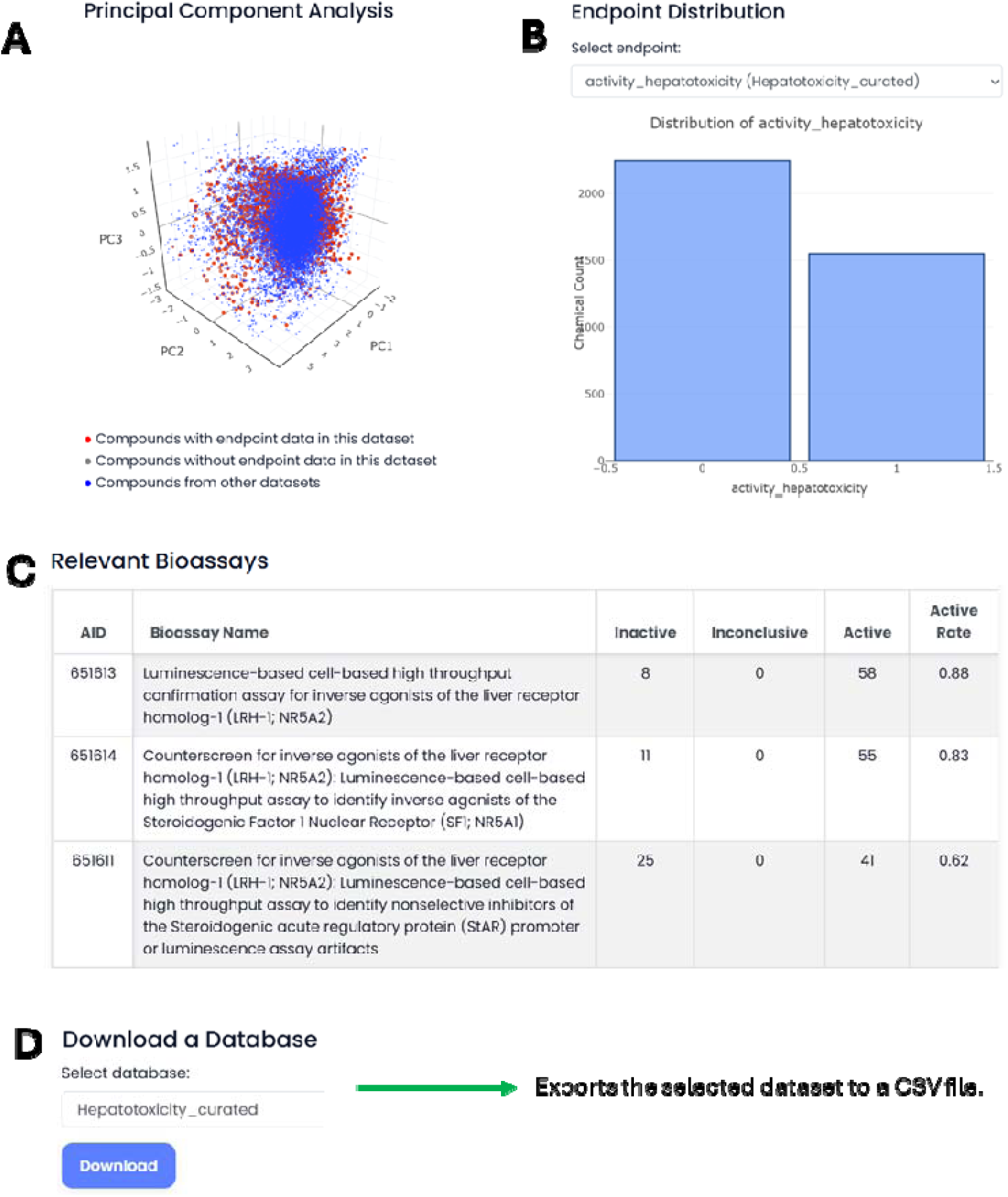
Overview of the Database functions. (A) Chemical space visualization. (B) Endpoint distribution. (C) Relevant bioassays for a selected endpoint. (D) Menu for downloading a selected dataset.

### Cheminformatics

The Cheminformatics module provides a modeling workflow of dataset upload, chemical curation, visualization, and QSAR modeling (Figure 6). The module allows users to efficiently generate machine learning models, compare their performance, and make predictions. For example, as shown in Figures 8, ToxiVerse provides a sample dataset as part of tutorial for user to learn the functions in the modeling and prediction workflow. This module enables researchers without programming expertise to carry out end-to-end QSAR modeling using ToxiVerse tools.

**Figure 6.**
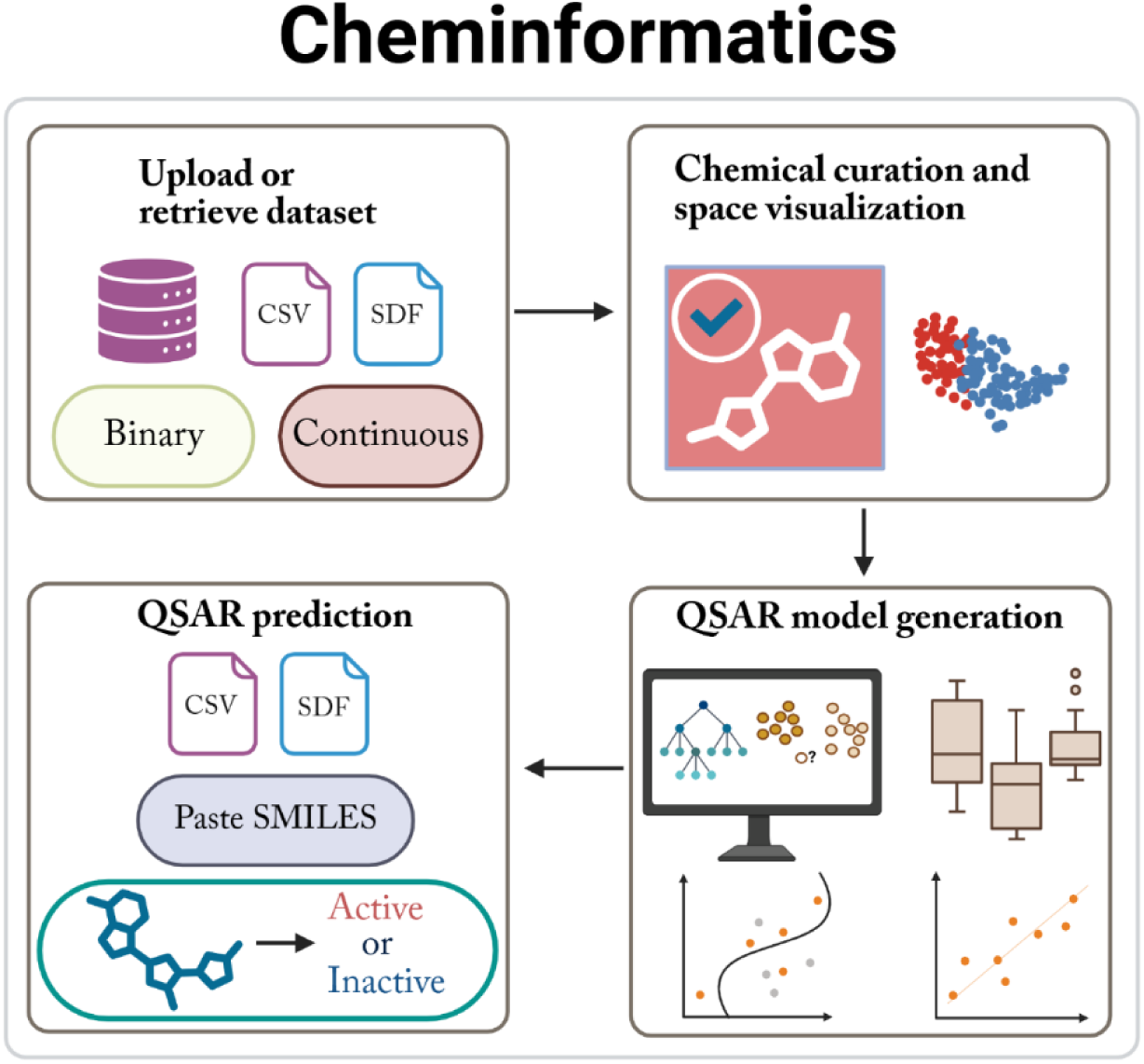
Functionalities of the Cheminformatics module.

### Dataset Upload and Retrieval

It enables users to upload datasets in SDF or CSV formats for computational modeling. The interface accepts data that includes user-defined column names for compound identifiers, SMILES, and activity values, along with the dataset type (binary or continuous). The uploaded dataset displayed in the interface can be downloaded as a CSV file using the *Download Dataset as CSV* option or removed using the *Remove Dataset* option (Figure 7A). Datasets can also be imported from PubChem using associated AIDs. The chemical-bioactivity records are retrieved initially for a given AID and activity outcomes are standardized by discarding *inconclusive* or *unspecified* results. If duplicate compound entries are present, the entry with a defined activity value is retained and duplicates are removed. The cleaned dataset is merged with InChI representations.

**Figure 7.**
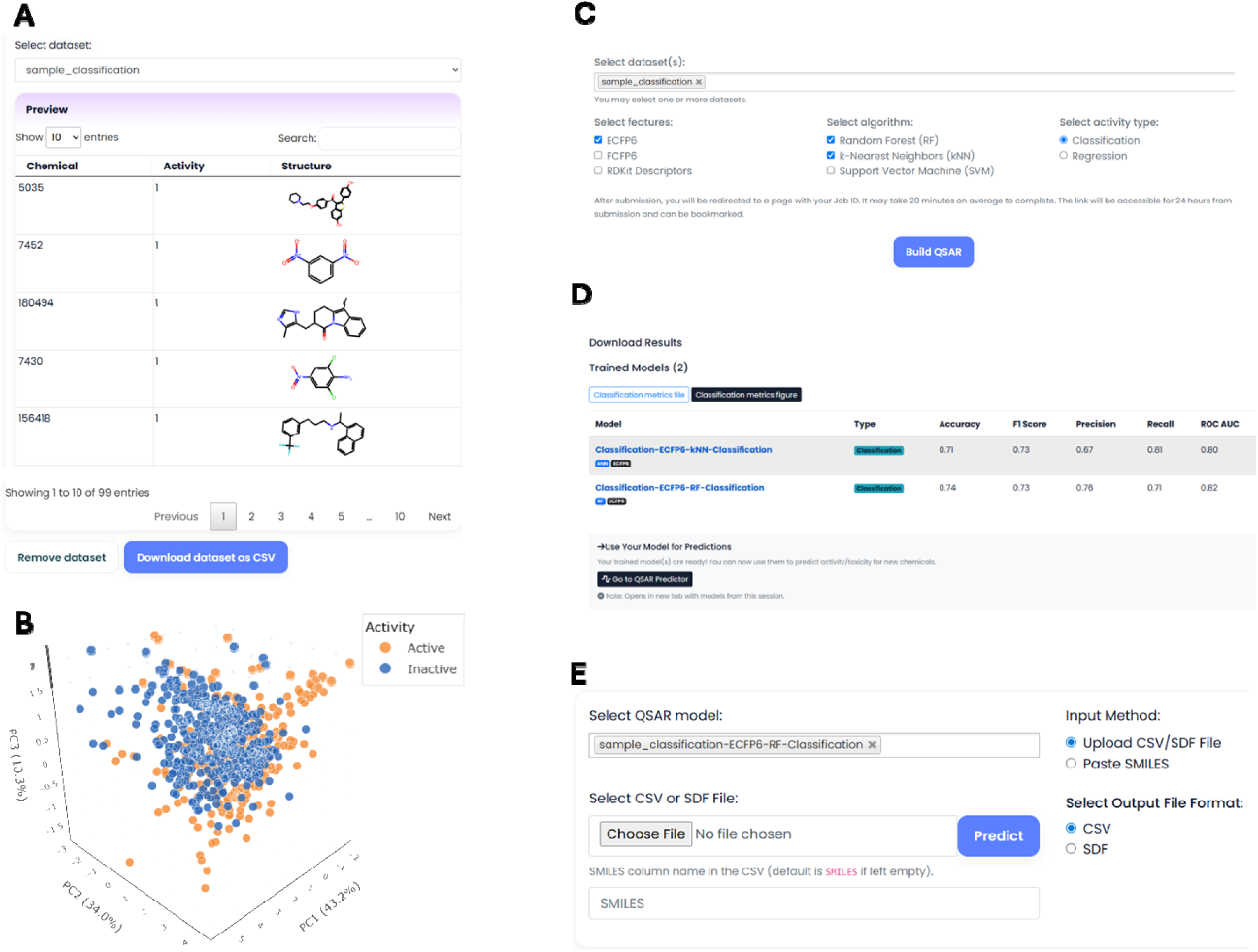
Example of main functions in Cheminformatics module using a sample classification dataset. (A) Display of the uploaded dataset in tabular format. (B) Chemical space visualization of the dataset. (C) Selection of an uploaded dataset for QSAR model generation. (D) Display of model metrics in tabular format and the link to access the generated models. (E) Prediction of new chemicals using a selected model.

### Structure Curation and Chemical Space Visualization

Chemical structure curation uses the ChEMBL Structure Pipeline with RDKit. This process identifies and corrects problematic structures, ensuring standardized, high-quality structures for modeling. This function also allows users to choose how duplicates should be handled, by retaining the highest or lowest activity, averaging activity values, or removing duplicates entirely. Users can choose to either overwrite the original dataset with the curated version or create a new dataset, in which case “_curated” will be added to the original dataset name. In the latter case, both the uploaded and curated datasets remain accessible. For chemical space visualization, PCA is performed on a set of molecular descriptors (calculated via RDKit). The resulting 3D plot, which uses the top three principal components as the coordinates, color-codes compounds by their activity value, allowing users to visually explore structural patterns or clusters in their dataset (Figure 7B).

### QSAR Model generation

The QSAR builder supports building ML models for toxicity datasets selected by users (Figure 7C and D). Molecular descriptors or fingerprints (Morgan/ECFP, FCFP, and RDKit descriptors) are computed and normalized when necessary. Classification and regression modeling using RF, SVM, and k-NN algorithms can be performed. Models are tuned using grid search with 5-fold cross-validation, and classification outputs use a probability threshold of 0.5 for assigning binary labels. The best-performing model is saved for future predictions. Performance metrics are accuracy, AUC, F1-score, precision, recall and ROC AUC for classification. Regression models are evaluated using R², Mean Squared Error (MSE), and Mean Absolute Percentage Error (MAPE). For example, we modeled a toxicity dataset (Supplementary File S1) of 888 compounds (444 actives, 444 inactives) (28) from a PubChem estrogen receptor antagonist assay (AID 1259248). Nine classification models (combinations of ECFP6, FCFP6, RDKit descriptors with kNN, RF, SVM) were built and evaluated via 5-fold cross-validation. Their performance metrics are shown in Supplementary Figure S2. The QSAR builder option allows users to download performance metrics plot for their models, along with the corresponding performance scores in a CSV file (Figure 7D). The results can be accessed for up to 24 hours from job submission via unique job link.

### QSAR Prediction

The QSAR prediction interface allows users to predict activities/toxicities for chemicals using trained models (Figure 7E). Chemicals to be predicted can be provided via file upload (CSV or SDF) or by direct SMILES input. Invalid structures are automatically removed. For CSV uploads, the system automatically detects a column named “SMILES,” or users can manually specify the SMILES column name. A “Paste Sample SMILES” feature is provided to help users quickly make predictions without uploading files. Predicted outputs are appended to the original dataset with model-specific columns named *ModelName_Prediction*. The results can be downloaded as CSV or SDF files.

### Distinctive Features of ToxiVerse

Despite the valuable contributions of other existing tools, most rely on fixed, pre-trained models for a predetermined set of endpoints, offering little to no flexibility for users to input new datasets or build customized predictive models (11). This rigidity restricts their utility where new data or endpoints need to be modeled and evaluated. Additionally, batch processing capabilities are often lacking. While many platforms are web-accessible, their functionalities can still pose barriers for users without programming backgrounds. ToxiVerse addresses these challenges through a fully integrated, web-based architecture that enables users to build custom QSAR models using either curated ToxiVerse datasets or their own experimental data. Its Bioprofiler module enriches chemical descriptors by leveraging high-throughput bioactivity data from PubChem and incorporates machine learning-based imputation to fill data gaps of selected assays. The platform also provides access to a rigorously curated database for various toxicity endpoints. Furthermore, ToxiVerse supports batch processing and step-by-step model building without requiring coding skills. By combining flexibility, mechanistic insight, and user accessibility in a single platform, ToxiVerse fills some of the critical usability and adaptability gaps left by earlier toxicity prediction tools.

## ACKNOWLEDGEMENT

This work was partially supported by the National Institute of Child Health and Human Development (Grant UC2HD113039) and the National Institute of Environmental Health Sciences (Grants R01ES031080 and R35ES031709).

---

**Supplementary Figure S1.** The Bioprofiler functions overview: generating five distinct outputs, each available for download with sample outputs provided. (A) Heatmap representation of initial bioprofile. (B) Box plot of performance metrics for models built on the selected assays. (C) The initial bioprofile shown as chemical-bioactivity matrix with a tabular format of compound activity outcomes (-1 = inactive, 0 = inconclusive, 1 = active). (D) The performance metrics file containing evaluation results for the models based on selected assays. (E) A complete bioprofile with filled data gaps. A represents Assay and C represents chemical in the figure.

**Supplementary Figure S2.** Performance metrics of the classification models for the estrogen receptor dataset.

**Supplementary File S1**: Toxicity dataset of 888 compounds (444 actives, 444 inactives) from a PubChem estrogen receptor antagonist assay (AID 1259248).

## REFERENCES

1. Guo W, Liu J, Dong F, Song M, Li Z, Khan MKH, et al. Review of machine learning and deep learning models for toxicity prediction. Exp Biol Med. 2023 Dec 6;15353702231209421.

2. Muster W, Breidenbach A, Fischer H, Kirchner S, Müller L, Pähler A. Computational toxicology in drug development. Drug Discov Today. 2008 Apr;13(7–8):303–10.

3. Waring MJ, Arrowsmith J, Leach AR, Leeson PD, Mandrell S, Owen RM, et al. An analysis of the attrition of drug candidates from four major pharmaceutical companies. Nat Rev Drug Discov. 2015 July;14(7):475–86.

4. Ford KA. Refinement, Reduction, and Replacement of Animal Toxicity Tests by Computational Methods. ILAR J. 2016 Dec;57(2):226–33.

5. Pognan F, Beilmann M, Boonen HCM, Czich A, Dear G, Hewitt P, et al. The evolving role of investigative toxicology in the pharmaceutical industry. Nat Rev Drug Discov. 2023 Apr;22(4):317–35.

6. Gibb S. Toxicity testing in the 21st century: A vision and a strategy. Reprod Toxicol. 2008 Jan;25(1):136–8.

7. Wang N, Li X, Xiao J, Liu S, Cao D. Data-driven toxicity prediction in drug discovery: Current status and future directions. Drug Discov Today. 2024 Nov;29(11):104195.

8. Shen J, Nicolaou CA. Molecular property prediction: recent trends in the era of artificial intelligence. Drug Discov Today Technol. 2019 Dec;32–33:29–36.

9. Jia X, Wang T, Zhu H. Advancing Computational Toxicology by Interpretable Machine Learning. Environ Sci Technol. 2023 Nov 21;57(46):17690–706.

10. Richard AM, Huang R, Waidyanatha S, Shinn P, Collins BJ, Thillainadarajah I, et al. The Tox21 10K Compound Library: Collaborative Chemistry Advancing Toxicology. Chem Res Toxicol. 2021 Feb 15;34(2):189–216.

11. Seal S, Mahale M, García-Ortegón M, Joshi CK, Hosseini-Gerami L, Beatson A, et al. Machine Learning for Toxicity Prediction Using Chemical Structures: Pillars for Success in the Real World. Chem Res Toxicol. 2025 May 19;38(5):759–807.

12. Fu L, Shi S, Yi J, Wang N, He Y, Wu Z, et al. ADMETlab 3.0: an updated comprehensive online ADMET prediction platform enhanced with broader coverage, improved performance, API functionality and decision support. Nucleic Acids Res. 2024 July 5;52(W1):W422–31.

13. Gu Y, Yu Z, Wang Y, Chen L, Lou C, Yang C, et al. admetSAR3.0: a comprehensive platform for exploration, prediction and optimization of chemical ADMET properties. Nucleic Acids Res. 2024 July 5;52(W1):W432–8.

14. Banerjee P, Kemmler E, Dunkel M, Preissner R. ProTox 3.0: a webserver for the prediction of toxicity of chemicals. Nucleic Acids Res. 2024 July 5;52(W1):W513–20.

15. Ji C, Svensson F, Zoufir A, Bender A. eMolTox: prediction of molecular toxicity with confidence. Valencia A, editor. Bioinformatics. 2018 July 15;34(14):2508–9.

16. Di Stefano M, Galati S, Piazza L, Granchi C, Mancini S, Fratini F, et al. VenomPred 2.0: A Novel *In Silico* Platform for an Extended and Human Interpretable Toxicological Profiling of Small Molecules. J Chem Inf Model. 2024 Apr 8;64(7):2275–89.

17. Braga RC, Alves VM, Silva MFB, Muratov E, Fourches D, Lião LM, et al. Pred-hERG: A Novel web-Accessible Computational Tool for Predicting Cardiac Toxicity. Mol Inform. 2015 Oct;34(10):698–701.

18. Borba JVB, Braga RC, Alves VM, Muratov EN, Kleinstreuer N, Tropsha A, et al. Pred-Skin: A Web Portal for Accurate Prediction of Human Skin Sensitizers. Chem Res Toxicol. 2021 Feb 15;34(2):258–67.

19. Lee HM, Yu MS, Kazmi SR, Oh SY, Rhee KH, Bae MA, et al. Computational determination of hERG-related cardiotoxicity of drug candidates. BMC Bioinformatics. 2019 May;20(S10):250.

20. Ruusmann V, Sild S, Maran U. QSAR DataBank repository: open and linked qualitative and quantitative structure–activity relationship models. J Cheminformatics. 2015 Dec;7(1):32.

21. Benfenati E, Manganaro A, Gini G. VEGA-QSAR: AI inside a platform for predictive toxicology.

22. Schultz TW, Diderich R, Kuseva CD, Mekenyan OG. The OECD QSAR Toolbox Starts Its Second Decade. In: Nicolotti O, editor. Computational Toxicology [Internet]. New York, NY: Springer New York; 2018 [cited 2025 Sept 1]. p. 55–77. (Methods in Molecular Biology; vol. 1800). Available from: http://link.springer.com/10.1007/978-1-4939-7899-1_2

23. Wu L, Yan B, Han J, Li R, Xiao J, He S, et al. TOXRIC: a comprehensive database of toxicological data and benchmarks. Nucleic Acids Res. 2023 Jan 6;51(D1):D1432–45.

24. Yang H, Sun L, Li W, Liu G, Tang Y. In Silico Prediction of Chemical Toxicity for Drug Design Using Machine Learning Methods and Structural Alerts. Front Chem. 2018 Feb 20;6:30.

25. Russo DP, Kim MT, Wang W, Pinolini D, Shende S, Strickland J, et al. CIIPro: a new read-across portal to fill data gaps using public large-scale chemical and biological data. Wren J, editor. Bioinformatics. 2017 Feb 1;33(3):464–6.

26. Zhu H. Supporting read-across using biological data. ALTEX. 2016;167–82.

27. Low Y, Sedykh A, Fourches D, Golbraikh A, Whelan M, Rusyn I, et al. Integrative Chemical–Biological Read-Across Approach for Chemical Hazard Classification. Chem Res Toxicol. 2013 Aug 19;26(8):1199–208.

28. 28. Ciallella HL, Chung E, Russo DP, Zhu H. Automatic Quantitative Structure–Activity Relationship Modeling to Fill Data Gaps in High-Throughput Screening. In: Zhu H, Xia M, editors. High-Throughput Screening Assays in Toxicology [Internet]. New York, NY: Springer US; 2022 [cited 2025 June 28]. p. 169–87. (Methods in Molecular Biology; vol. 2474). Available from: https://link.springer.com/10.1007/978-1-0716-2213-1_16

29. Kim S, Chen J, Cheng T, Gindulyte A, He J, He S, et al. PubChem 2025 update. Nucleic Acids Res. 2025 Jan 6;53(D1):D1516–25.

30. Breiman L. Random Forests. Mach Learn. 2001 Oct;45(1):5–32.

31. Rogers D, Hahn M. Extended-Connectivity Fingerprints. J Chem Inf Model. 2010 May 24;50(5):742–54.

32. Wang W, Kim MT, Sedykh A, Zhu H. Developing Enhanced Blood–Brain Barrier Permeability Models: Integrating External Bio-Assay Data in QSAR Modeling. Pharm Res. 2015 Sept;32(9):3055–65.

33. Zhao L, Wang W, Sedykh A, Zhu H. Experimental Errors in QSAR Modeling Sets: What We Can Do and What We Cannot Do. ACS Omega. 2017 June 30;2(6):2805–12.

34. Kim MT, Sedykh A, Chakravarti SK, Saiakhov RD, Zhu H. Critical Evaluation of Human Oral Bioavailability for Pharmaceutical Drugs by Using Various Cheminformatics Approaches. Pharm Res. 2014 Apr;31(4):1002–14.

35. Zhao L, Wang W, Sedykh A, Zhu H. Experimental Errors in QSAR Modeling Sets: What We Can Do and What We Cannot Do. ACS Omega. 2017 June 30;2(6):2805–12.

36. Chung E, Russo DP, Ciallella HL, Wang YT, Wu M, Aleksunes LM, et al. Data-Driven Quantitative Structure–Activity Relationship Modeling for Human Carcinogenicity by Chronic Oral Exposure. Environ Sci Technol. 2023 Apr 25;57(16):6573–88.

37. Ciallella HL, Russo DP, Sharma S, Li Y, Sloter E, Sweet L, et al. Predicting Prenatal Developmental Toxicity Based On the Combination of Chemical Structures and Biological Data. Environ Sci Technol. 2022 May 3;56(9):5984–98.

38. Aljarf R, Tang S, Pires DEV, Ascher DB. embryoTox: Using Graph-Based Signatures to Predict the Teratogenicity of Small Molecules. J Chem Inf Model. 2023 Jan 23;63(2):432–41.

39. Knox C, Wilson M, Klinger CM, Franklin M, Oler E, Wilson A, et al. DrugBank 6.0: the DrugBank Knowledgebase for 2024. Nucleic Acids Res. 2024 Jan 5;52(D1):D1265–75.

40. Klopman G, Stuart SE. Multiple computer-automated structure evaluation study of aquatic toxicity. III. Vibrio fischeri. Environ Toxicol Chem. 2003 Mar 1;22(3):466–72.

41. Mulliner D, Schmidt F, Stolte M, Spirkl HP, Czich A, Amberg A. Computational Models for Human and Animal Hepatotoxicity with a Global Application Scope. Chem Res Toxicol. 2016 May 16;29(5):757–67.

42. Zhu H, Ye L, Richard A, Golbraikh A, Wright FA, Rusyn I, et al. A Novel Two-Step Hierarchical Quantitative Structure–Activity Relationship Modeling Work Flow for Predicting Acute Toxicity of Chemicals in Rodents. Environ Health Perspect. 2009 Aug;117(8):1257–64.

43. Sedykh A, Fourches D, Duan J, Hucke O, Garneau M, Zhu H, et al. Human Intestinal Transporter Database: QSAR Modeling and Virtual Profiling of Drug Uptake, Efflux and Interactions. Pharm Res. 2013 Apr;30(4):996–1007.

44. Moda TL, Montanari CA, Andricopulo AD. Hologram QSAR model for the prediction of human oral bioavailability. Bioorg Med Chem. 2007 Dec;15(24):7738–45.

45. Ciallella HL, Russo DP, Aleksunes LM, Grimm FA, Zhu H. Predictive modeling of estrogen receptor agonism, antagonism, and binding activities using machine- and deep-learning approaches. Lab Invest. 2021 Apr;101(4):490–502.

46. Bento AP, Hersey A, Félix E, Landrum G, Gaulton A, Atkinson F, et al. An open source chemical structure curation pipeline using RDKit. J Cheminformatics. 2020 Dec;12(1):51.

47. Cortes C, Vapnik V. Support-vector networks. Mach Learn. 1995 Sept;20(3):273–97.

48. Cover T, Hart P. Nearest neighbor pattern classification. IEEE Trans Inf Theory. 1967 Jan;13(1):21–7.

49. Daood NJ, Russo DP, Chung E, Qin X, Zhu H. Predicting Chemical Immunotoxicity through Data-Driven QSAR Modeling of Aryl Hydrocarbon Receptor Agonism and Related Toxicity Mechanisms. Environ Health. 2024 July 19;2(7):474–85.

50. Chung E, Wen X, Jia X, Ciallella HL, Aleksunes LM, Zhu H. Hybrid non-animal modeling: A mechanistic approach to predict chemical hepatotoxicity. J Hazard Mater. 2024 June;471:134297.

51. Jia X, Wen X, Russo DP, Aleksunes LM, Zhu H. Mechanism-driven modeling of chemical hepatotoxicity using structural alerts and an in vitro screening assay. J Hazard Mater. 2022 Aug;436:129193.

