## Supplementary Figure S1 for "ToxiVerse: A Public Platform for Chemical Toxicity Data Sharing and Customizable Predictive Modeling"

**A**

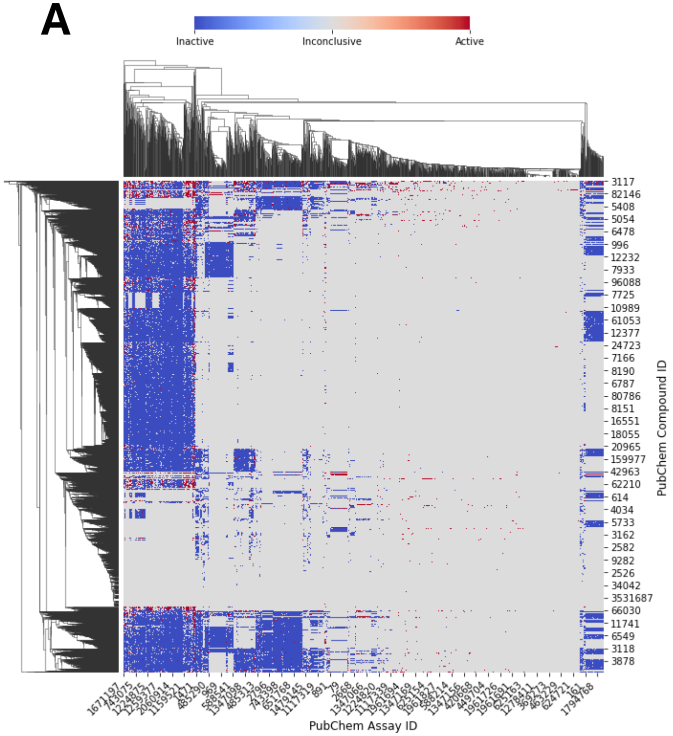


Heatmap

**B**

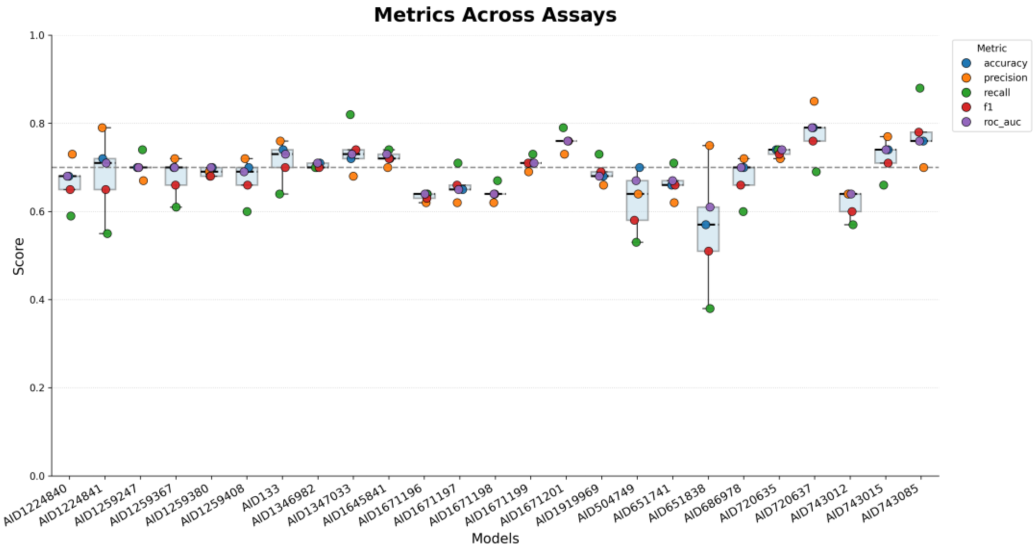


Metrics Plot

**C**

Initial Bioprofile

| CID | A1 | A2 | A3 | A4 | ... | An |
|---|---|---|---|---|---|---|
| C1 | 1 | -1 | 0 | 0 | ... | -1 |
| C2 | -1 | 0 | 0 | -1 | ... | 0 |
| C3 | 0 | 0 | 0 | 0 | ... | 0 |
| C4 | 1 | 0 | -1 | 0 | ... | 1 |
| ... | ... | ... | ... | ... | ... | ... |
| Cn | -1 | 0 | 0 | 0 | ... | 0 |

**D**

Model Metrics

| Model | accuracy | precision | recall | f1 | roc_auc |
|---|---|---|---|---|---|
| A1 | 0.68 | 0.73 | 0.59 | 0.65 | 0.68 |
| A2 | 0.72 | 0.79 | 0.55 | 0.65 | 0.71 |
| A3 | 0.7 | 0.68 | 0.7 | 0.69 | 0.7 |
| A4 | 0.7 | 0.67 | 0.74 | 0.7 | 0.7 |
| ... | ... | ... | ... | ... | ... |
| A25 | 0.7 | 0.72 | 0.61 | 0.66 | 0.7 |

**E**

Complete Bioprofile

| CID | A1 | A2 | A3 | A4 | ... | An |
|---|---|---|---|---|---|---|
| C1 | 1 | -1 | -1 | 1 | ... | -1 |
| C2 | -1 | -1 | -1 | -1 | ... | -1 |
| C3 | -1 | 1 | -1 | -1 | ... | -1 |
| C4 | 1 | -1 | -1 | -1 | ... | 1 |
| ... | ... | ... | ... | ... | ... | ... |
| Cn | -1 | 1 | 1 | -1 | ... | -1 |
