## Supplementary figures and images for "ToxiVerse: A Public Platform for Chemical Toxicity Data Sharing and Customizable Predictive Modeling"

### Supplementary Figure S2

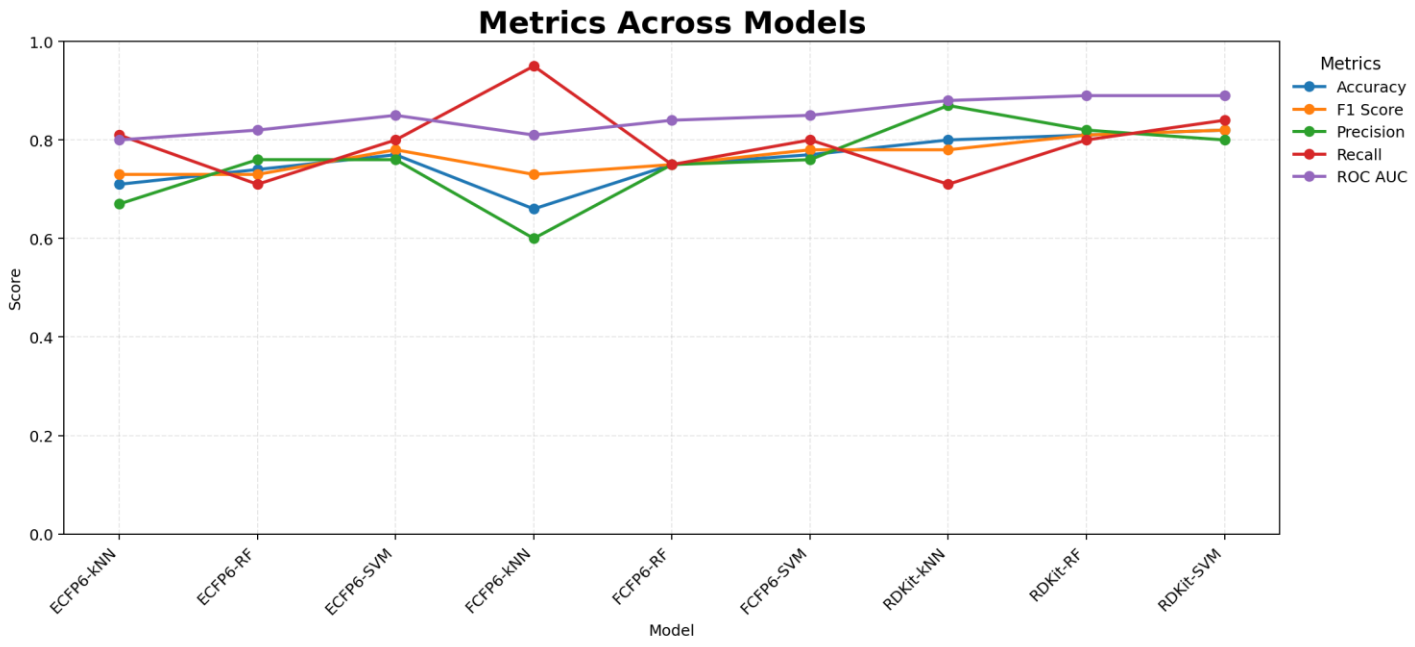
